# High-titer convalescent serum from cats recovered from severe fever with thrombocytopenia syndrome protects SFTSV-infected mice

**DOI:** 10.64898/2026.10.07.756134

**Authors:** Hirohisa Mekata, Mari Yamamoto, Takeshi Kawaguchi, Tamaki Okabayashi

**Affiliations:** Center for Animal Disease Control, University of Miyazaki, 1-1 Gakuen-Kibanadai-Nishi, Miyazaki 889-2192, Japan; Division of Respirology, Rheumatology, Infectious Diseases, and Neurology, Department of Internal Medicine, Faculty of Medicine, University of Miyazaki, 5200 Kihara, Kiyotake, Miyazaki 889-1692, Japan

**Keywords:** severe fever with thrombocytopenia syndrome, cats, convalescent serum, neutralizing antibody, passive immunotherapy

## Abstract

Severe fever with thrombocytopenia syndrome (SFTS) is an emerging, tick-borne zoonosis caused by severe fever with thrombocytopenia syndrome virus (SFTSV) and is associated with high case-fatality rates in humans and cats. No vaccines or antiviral drugs are currently available for the prevention or treatment of feline SFTS. Because cats with SFTS often develop blood viral RNA loads exceeding those in severe human cases, we hypothesized that some naturally recovered cats develop strong neutralizing antibody responses and may serve as a potential source of antibody-rich serum for passive immunotherapy. We compared 80% plaque-reduction neutralization test (PRNT_80_) titers in convalescent sera from six human SFTS survivors and 14 cats that had recovered from naturally acquired SFTS, then evaluated the therapeutic efficacy of convalescent cat serum in SFTSV-infected, interferon receptor-deficient AG129 mice. Some SFTS-recovered cats showed high PRNT_80_ titers. Passive administration of high-titer convalescent cat serum early after infection protected SFTSV-infected mice. A single dose given at 2 or 3 days post-infection completely protected mice, whereas treatment at 4 days post-infection resulted in 40% survival. Administration on two consecutive days was associated with lower late-phase viremia in surviving mice. Sera from three independent SFTS-recovered cats improved survival, with greater effects observed for sera with higher neutralizing antibody titers. These findings indicate that some cats recovering from SFTS can develop high neutralizing antibody titers and their sera can exert therapeutic effects in an SFTSV-infected mouse model. These findings support further evaluation of passive immunotherapy as a treatment strategy for feline SFTS.

## Introduction

Severe fever with thrombocytopenia syndrome (SFTS) is an emerging, potentially fatal tick-borne zoonosis caused by severe fever with thrombocytopenia syndrome virus (SFTSV), an enveloped, tripartite, negative-sense RNA virus classified within the species *Bandavirus dabieense*, genus *Bandavirus*, family *Phenuiviridae* (International Committee on Taxonomy of Viruses, 2024). Human SFTS is characterized by fever, gastrointestinal symptoms, leukopenia, thrombocytopenia, and multiorgan involvement (Kato et al., 2016). Since its first report in China in 2011 (Yu et al., 2011), SFTS has been identified in South Korea (Park et al., 2014), Japan (Takahashi et al., 2014), and several Southeast Asian countries (Tran et al., 2019; Rattanakomol et al., 2022), with reported case-fatality rates of approximately 5%–27% (Huang et al., 2021; Lee et al., 2024; Kobayashi et al., 2020).

Domestic cats are highly susceptible to SFTSV infection, with a reported case-fatality rate of 58%–63% (Matsuu et al., 2019; Ando et al., 2021). Although animal SFTS is not legally notifiable in Japan, 194 feline SFTS cases were reported through a domestic animal SFTS diagnostic network in 2024, exceeding the number of reported human SFTS cases in the same year (National Institute of Infectious Diseases, 2024). Direct cat-to-human transmission of SFTSV has also been documented among veterinary personnel and people caring for infected cats (Yamanaka et al., 2020; Mekata et al., 2023). However, no effective treatment or vaccine is currently available for feline SFTS.

Passive immunization with convalescent serum has been explored for acute viral infections (Libster et al., 2021; van Griensven et al., 2016); however, evidence for SFTS remains limited. In an SFTSV-infected mouse model, administration of serum from patients who had recovered from SFTS resulted in 20% survival when treatment was initiated 1 day after infection (Shimada et al., 2015). Because cats with SFTS often develop blood viral RNA loads exceeding those observed in severe human cases (Park et al., 2019; Gai et al., 2012), we hypothesized that some naturally recovered cats develop strong neutralizing antibody responses and may serve as a potential source of antibody-rich sera for passive immunotherapy.

To contextualize the neutralizing antibody responses of recovered cats, we compared their titers with those of human SFTS survivors, then evaluated the therapeutic efficacy of convalescent cat serum in an SFTSV-infected mouse model. Since blood transfusion from donor cats is already performed in small animal practice following compatibility testing (Taylor et al., 2021), we considered whether passive immunotherapy could be a clinically plausible approach for feline SFTS.

## Materials and Methods

### Ethics statement

This study was approved by the Research Ethics Review Committee of the Faculty of Medicine, University of Miyazaki (Approval No. O-1434) for the use of serum samples from convalescent patients with SFTS. Written informed consent was obtained from all participants. Serum samples from cats that had recovered from SFTS were collected at private veterinary clinics after obtaining informed consent from their owners. All animal experimental protocols, including SFTSV infection studies using AG129 mice and the collection and use of serum samples from SFTS-recovered cats, were approved by the Institutional Animal Care and Use Committee of the University of Miyazaki (Approval No. 2025-055). All experiments involving infectious SFTSV were conducted in animal biosafety level 3 containment laboratories at the Center for Animal Disease Control, University of Miyazaki.

### Serum samples

Human sera were obtained from six convalescent patients who had been hospitalized at the University of Miyazaki Hospital with SFTS and subsequently recovered. SFTSV infection was confirmed using routine diagnostic testing performed at a prefectural institute of public health. Serum samples were collected during follow-up visits in the convalescent phase, 106–253 days after disease onset. The recovered patients ranged in age from 55 to 83 years (median 71 years), and included two males and four females.

Cat serum samples were obtained from 14 cats that had been examined at private veterinary clinics and subsequently recovered from SFTS. SFTSV infection was confirmed in our laboratory using reverse transcription quantitative PCR (RT-qPCR) for detection of viral RNA. The recovered cats ranged in age from 1 to 12 years (median 3.5 years), and included five males and nine females. Sera were collected during follow-up visits in the convalescent phase, 31–690 days after disease onset. All convalescent cat serum samples used for administration to mice were confirmed to be negative for detectable SFTSV RNA by RT-qPCR before use.

### Cells and virus

The A17 SFTSV strain was originally isolated from a tick in Miyazaki Prefecture, Japan (Sato et al., 2021). For the neutralizing antibody assay, a cell culture-adapted A17 strain passaged 20 times was used because it formed clear plaques in the assay system. Vero cells stably expressing dendritic cell-specific intercellular adhesion molecule-3-grabbing non-integrin were used to enhance susceptibility to SFTSV (Mekata et al., 2025). In contrast, a tick-derived A17 strain without prior cell culture adaptation was used for animal experiments.

### Mice

AG129 mice, a genetically modified strain deficient in type I and type II interferon receptors, were used in this study. Male and female mice aged 8–10 weeks were used for infection experiments. They were obtained under a breeding and use license from B&K Universal Ltd. (East Yorkshire, UK), a Marshall BioResources company, and maintained as an in-house breeding colony.

### Neutralizing antibody assay

Neutralizing antibody titers in sera from convalescent patients with SFTS and SFTS-recovered cats were determined using an 80% plaque reduction neutralization test (PRNT_80_). Serum samples were heat-inactivated at 56 °C for 30 min and diluted in phosphate-buffered saline (PBS). Subsequently, sera were serially two-fold diluted from 1:20 to 1:10240 and mixed with an equal volume of virus adjusted to 500 plaque-forming units/mL. After incubation at 37 °C for 1 h, serum–virus mixtures were inoculated onto cell monolayers. Following incubation under a methylcellulose overlay, plaques were visualized by crystal violet staining and counted. The plaque reduction rate at each serum dilution was calculated relative to that in serum-free controls. The PRNT_80_ titer was defined as the serum dilution corresponding to an 80% reduction in plaque number and was calculated using a four-parameter logistic regression model in GraphPad Prism 6 (GraphPad Software, San Diego, CA, USA).

### Animal experiments

Five to six mice were used per group. Each mouse was subcutaneously inoculated with 200 μL of virus solution containing 200 focus-forming units of SFTSV strain A17. All serum samples used for administration were heat-inactivated at 56 °C for 30 min before use.

Therapeutic studies using serum from SFTS-recovered cats were performed in AG129 mice under three treatment regimens. In the first regimen, serum from a cat with a neutralizing antibody titer of PRNT_80_ = 1408 was administered intraperitoneally as a single 400 μL dose at 2, 3, or 4 dpi. In the second regimen, the same serum was administered intraperitoneally twice (400 μL per dose) at 2 and 3 dpi, 3 and 4 dpi, 4 and 5 dpi, or 5 and 6 dpi. In the third regimen, sera from three individual SFTS-recovered cats, distinct from the serum used in the first and second regimens, were administered intraperitoneally as a single 400 μL dose at 3 dpi. The PRNT_80_ titers of Cat #2, Cat #3, and Cat #4 sera were 1243, 315, and 617, respectively. In the first regimen, control mice received the same volume of SFTSV-seronegative and SFTSV RNA-negative healthy cat serum at 2 dpi. In the second regimen, control mice received PBS at 2 and 3 dpi. In the third regimen, control mice received PBS at 3 dpi.

Blood samples (20 μL per mouse) were collected via tail-vein puncture at 5 and 10 dpi to measure viral RNA levels. In the second regimen, blood samples were collected at 6 and 13 dpi. Body weight was recorded daily throughout the experimental period. Mice were monitored for clinical signs including hunched posture, ruffled fur, and decreased activity.

A humane endpoint was defined as loss of more than 20% of initial body weight, at which point mice were euthanized.

### RNA extraction and RT-qPCR

Whole blood samples (20 μL) were mixed with 180 μL of virus inactivation reagent, Prep Buffer A (Precision System Science, Chiba, Japan). Total RNA was extracted using a fully automated nucleic acid extraction system (MagLEAD, Precision System Science). SFTSV RNA was quantified, and the internal control gene was simultaneously detected using One-Step PrimeScript III RT-qPCR Mix (TaKaRa Bio, Kusatsu, Japan) and PrimeTime qPCR assays (Integrated DNA Technologies, Coralville, IA, USA) with gene-specific primers and probes.

For quantification, a standard curve was generated using synthetic double-stranded DNA corresponding to a partial sequence of the SFTSV S segment (Integrated DNA Technologies). The DNA solution was adjusted to known concentrations, expressed as copies/μL, and serially diluted 10-fold using EASY Dilution (for Real Time PCR) (TaKaRa Bio). These standards were included in each RT-qPCR run. Values below the lower limit of quantification were assigned a value of 1,000 copies/mL for plotting and statistical analysis.

The primer and probe set used for quantification of SFTSV RNA was as follows: forward primer, TGTCAGAGTGGTCCAGGATT; reverse primer, ACCTGTCTCCTTCAGCTTCT; and probe, FAM-TGGAGTTTG/ZEN/GTGAGCAGCAGC-Iowa Black FQ (Yoshikawa et al., 2014). The primer and probe set used for detection of glyceraldehyde-3-phosphate dehydrogenase mRNA was as follows: forward primer, GCTGCCCAGAACATCATCC; reverse primer, GTCAGATCCACRACBRAYAC; and probe, ROX-TCACTGGCATGGCCTTCCGT-BHQ2 (Taglinger et al., 2008).

### Statistical analysis

Differences in PRNT_80_ titers between convalescent patients with SFTS and SFTS-recovered cats were analyzed using the Mann–Whitney U test. Survival curves were analyzed using the Kaplan–Meier method and compared using the log-rank (Mantel–Cox) test. Body weight data were analyzed using two-way repeated-measures analysis of variance followed by Dunnett’s multiple comparisons test. To avoid bias due to missing values caused by death or euthanasia at the humane endpoint, the analysis was restricted to time points before missing body-weight values occurred. Viral RNA levels at each time point were compared using the Kruskal–Wallis test followed by Dunn’s multiple comparisons test. *P* values <0.05 were considered statistically significant. All statistical analyses were performed using GraphPad Prism 6.

## Results

### Neutralizing antibody titers in SFTS-recovered cats and convalescent patients

To compare neutralizing antibody responses between SFTS-recovered cats and convalescent human patients, we measured PRNT_80_ titers in serum samples from six recovered patients and 14 recovered cats. The median PRNT_80_ titer was 937 (interquartile range, 428.2–2044) in convalescent cats and 404 (interquartile range, 180.3–689.5) in convalescent human patients (Figure 1). In this exploratory interspecies comparison, PRNT_80_ titers tended to be higher in cats than in humans, although the difference was not statistically significant (Mann–Whitney U test, *P* = 0.06).

**Figure 1.**
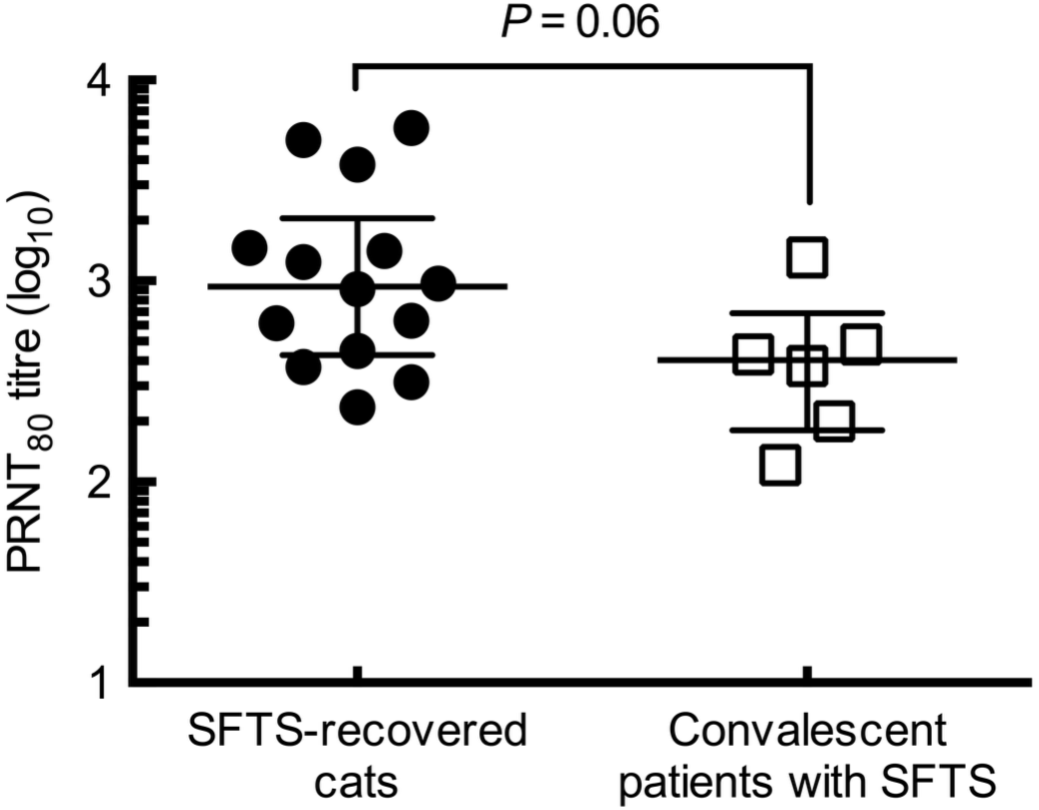
Neutralizing antibody titers in SFTS-recovered cats and convalescent patients with SFTS. Titers were measured using an 80% plaque reduction neutralization test (PRNT_80_) in serum samples from convalescent patients with severe fever with thrombocytopenia syndrome (SFTS) (*n* = 6) and SFTS-recovered cats (*n* = 14). Each dot represents an individual serum sample. Horizontal lines indicate medians and interquartile ranges. Groups were compared using the Mann–Whitney U test (*P* = 0.06).

#### Single-dose administration of convalescent cat serum protects SFTSV-infected mice

Because some convalescent cats exhibited high neutralizing antibody titers, we next investigated whether their sera could be used for passive immunotherapy in a murine model. SFTSV-infected, interferon receptor-deficient AG129 mice were administered convalescent cat serum (PRNT_80_ = 1408) at 2, 3, or 4 days post-infection (dpi). Control mice received serum from healthy cats with no history of SFTSV infection at 2 dpi. The log-rank test showed a significant difference in survival among groups (*P* = 0.0028) (Figure 2A). All control mice died by 8 dpi, resulting in 0% survival (0/5), whereas treatment at 2 or 3 dpi resulted in 100% survival (5/5) and treatment at 4 dpi resulted in 40% survival (2/5). Body weight loss was significantly attenuated in mice treated at 2 or 3 dpi but not in those treated at 4 dpi (Figure 2B). Earlier treatment tended to reduce blood viral RNA levels, with a significant reduction observed at 5 dpi in mice treated at 2 dpi (Figure 2C). However, viral RNA remained detectable in survivors at 10 dpi, indicating incomplete viral suppression after a single dose. These results indicate that single-dose convalescent cat serum provides therapeutic benefit when administered early after infection.

**Figure 2.**
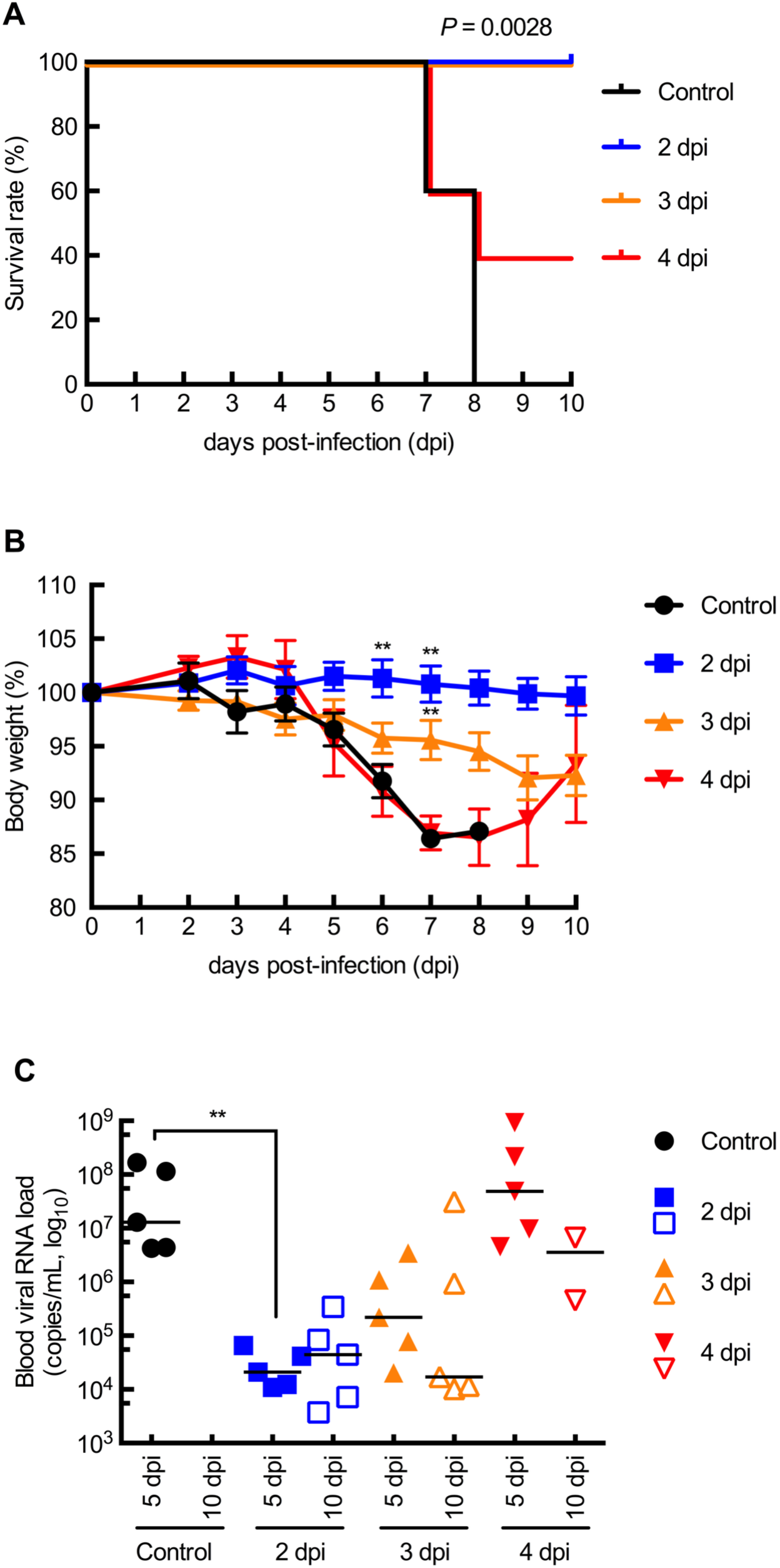
Single-dose administration of convalescent cat serum protects SFTSV-infected mice. AG129 mice were subcutaneously inoculated with 200 focus-forming units of the A17 strain of SFTSV and intraperitoneally administered a single dose of convalescent cat serum at 2, 3, or 4 days post-infection (dpi). Control mice received serum from healthy cats at 2 dpi. (A) Survival was analyzed using the Kaplan–Meier method and compared using the log-rank test. The *P* value indicates the overall log-rank test across groups. (B) Body weight changes are presented as the mean ± standard error of the mean. Data were analyzed using two-way repeated-measures analysis of variance (ANOVA) followed by Dunnett’s multiple comparisons test using data up to 7 dpi, when all animals were alive. (C) Blood viral RNA levels were measured using reverse transcription quantitative PCR. Each dot represents an individual mouse, and bars indicate medians. Filled and open symbols indicate samples collected at earlier and later time points, respectively. Data were analyzed using the Kruskal–Wallis test followed by Dunn’s multiple comparisons test. Asterisks indicate significant differences versus controls. Blood samples at 10 dpi were available only from surviving mice. \**P* < 0.05 and \*\**P* < 0.01.

### Two consecutive daily doses improve suppression of late-phase viremia

To determine whether therapeutic efficacy could be further improved, we evaluated administration of convalescent cat serum on two consecutive days. AG129 mice were infected with SFTSV and treated with convalescent cat serum at 2 and 3, 3 and 4, 4 and 5, or 5 and 6 dpi. Control mice received PBS injections at 2 and 3 dpi. The log-rank test showed a significant difference in survival among groups (*P* < 0.0001). All control mice died by 9 dpi, resulting in 0% survival (0/6) (Figure 3A). In contrast, mice treated at 2 and 3 dpi or 3 and 4 dpi were completely protected, with 100% survival (6/6). Mice treated at 3 and 4 dpi showed transient body weight loss exceeding 10% followed by subsequent recovery (Figure 3B). In contrast, treatment at 4 and 5 dpi or 5 and 6 dpi resulted in survival rates of 33% (2/6) and 0% (0/6), respectively. Administration on two consecutive days markedly reduced blood viral RNA levels at 13 dpi, suggesting enhanced suppression of late-phase viremia (Figure 3C).

**Figure 3.**
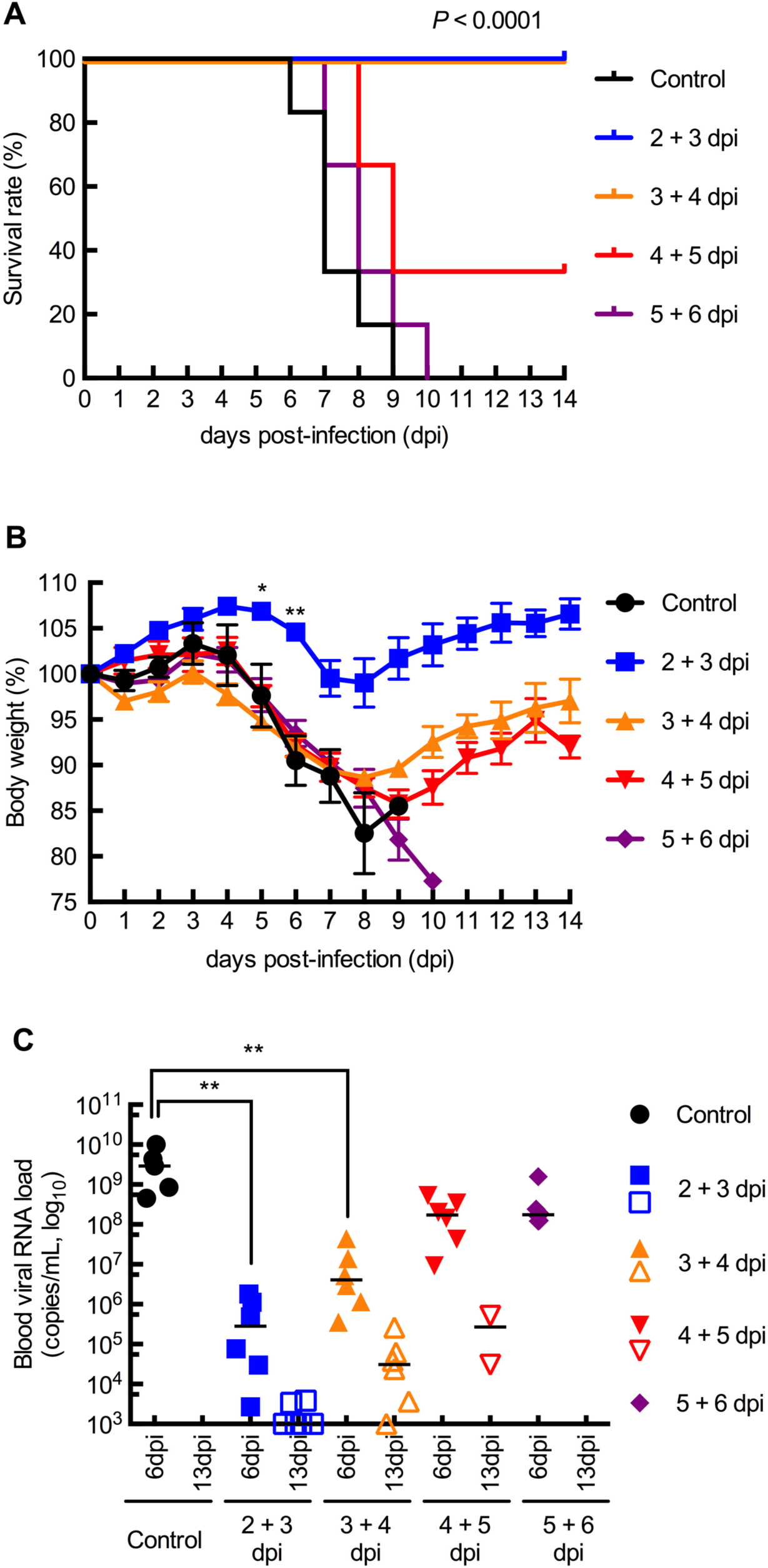
Therapeutic effect of administering convalescent cat serum on two consecutive days in SFTSV-infected mice. AG129 mice were subcutaneously inoculated with 200 focus-forming units of the A17 strain of SFTSV. Convalescent cat serum was administered intraperitoneally at 2 and 3, 3 and 4, 4 and 5, or 5 and 6 days post-infection (dpi). Control mice received phosphate-buffered saline at 2 and 3 dpi. (A) Survival was analyzed using the Kaplan–Meier method and compared using the log-rank test. The *P* value indicates the overall log-rank test across groups. (B) Body weight changes are presented as the mean ± standard error of the mean. Data were analyzed using two-way repeated-measures ANOVA followed by Dunnett’s multiple comparisons test for all time points at which all animals were alive. (C) Blood viral RNA levels were measured using RT-qPCR. Each dot represents an individual mouse, and bars indicate medians. Filled and open symbols indicate samples collected at earlier and later time points, respectively. Values below the lower limit of quantification were plotted at 1,000 copies/mL. Data were analyzed using the Kruskal–Wallis test followed by Dunn’s multiple comparisons test. Asterisks indicate significant differences versus controls. Blood samples at 6 and 13 dpi were available only from surviving mice. \**P* < 0.05 and \*\**P* < 0.01.

### Sera from individual convalescent cats confer therapeutic effects in SFTSV-infected mice

Finally, to determine whether therapeutic efficacy could be achieved using sera from other SFTS-recovered cats, we used sera from three individual SFTS-recovered cats with different neutralizing antibody titers. AG129 mice were infected with SFTSV and treated at 3 dpi with convalescent cat serum from the three individual cats: Cat #2, PRNT_80_ = 1243; Cat #3, PRNT_80_ = 315; and Cat #4, PRNT_80_ = 617. Control mice received PBS at 3 dpi. The log-rank test showed a significant difference in survival among groups (*P* < 0.0001). All control mice died by 9 dpi, resulting in 0% survival (0/6) (Figure 4A). Mice treated with serum from the cat with the highest neutralizing antibody titer (Cat #2) showed 100% survival (6/6) and exhibited significantly less body weight loss than control mice at 7 and 8 dpi (Figure 4B). Although mice treated with Cat #3 or Cat #4 serum showed transient body weight loss comparable to that of controls, only one mouse died in each group, resulting in 83% survival (5/6) in each group, and the remaining animals recovered (Figure 4A,B). Blood viral RNA levels at 5 dpi were significantly lower in mice treated with Cat #2 or Cat #4 serum than in control mice (Figure 4C). Collectively, these results suggest that sera from individual SFTS-recovered cats exert therapeutic effects in SFTSV-infected mice, although the magnitude of efficacy may depend partly on donor neutralizing antibody titers.

**Figure 4.**
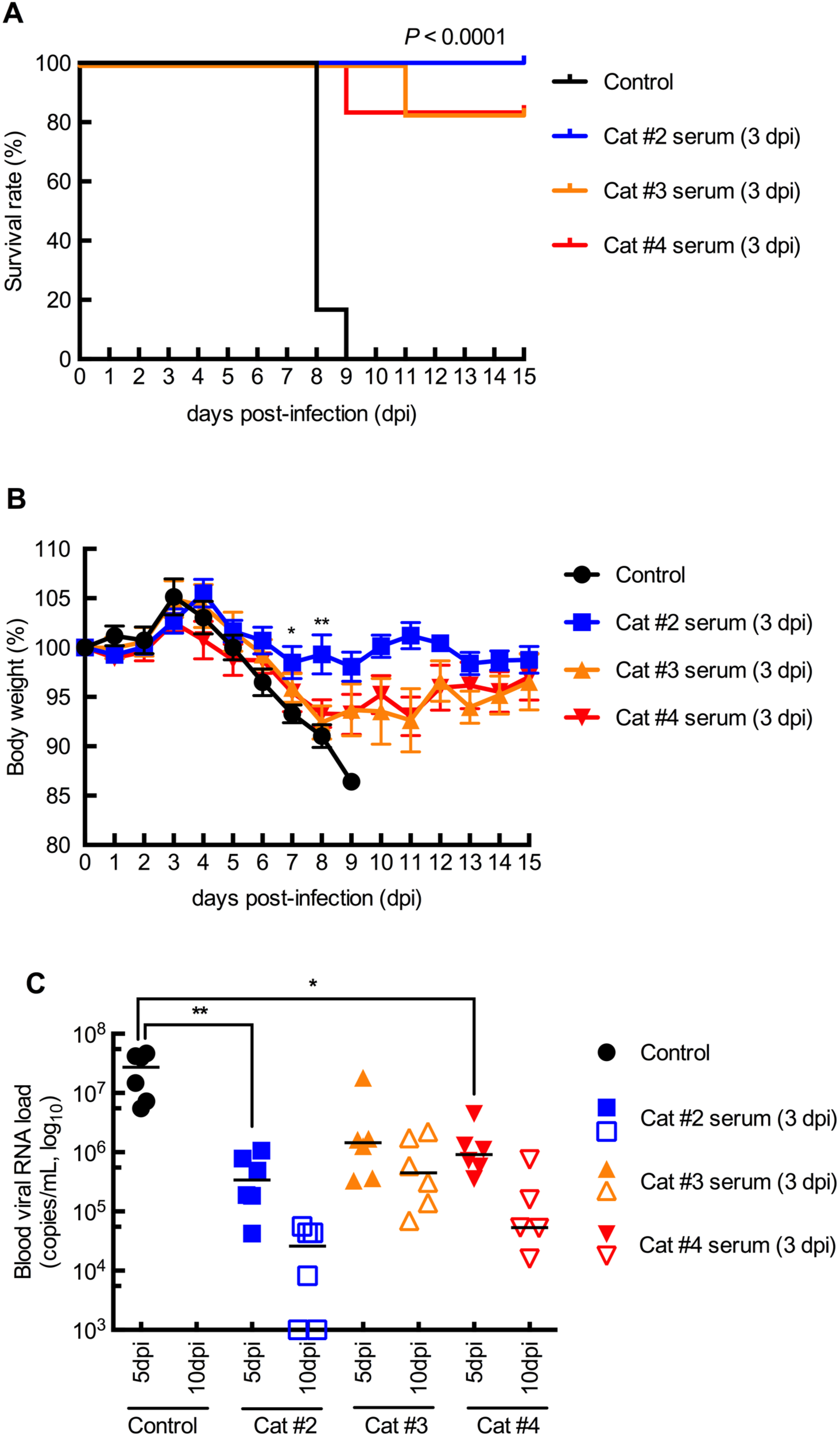
Sera from individual SFTS-recovered cats confer therapeutic effects in SFTSV-infected mice. AG129 mice were subcutaneously inoculated with 200 focus-forming units of the A17 strain of SFTSV and intraperitoneally administered a single dose of serum from one of the three SFTS-recovered cats at 3 dpi. Cat #2, Cat #3, and Cat #4 sera had PRNT_80_ titers of 1243, 315, and 617, respectively. Control mice received phosphate-buffered saline at 3 dpi. (A) Survival was analyzed using the Kaplan–Meier method and compared using the log-rank test. The *P* value indicates the overall log-rank test across groups. (B) Body weight changes are presented as the mean ± standard error of the mean. Data were analyzed using two-way repeated-measures ANOVA followed by Dunnett’s multiple comparisons test using data up to 8 dpi, as body-weight measurements were obtained before euthanasia based on the humane endpoint on that day. (C) Blood viral RNA levels were measured using RT-qPCR. Each dot represents an individual mouse and bars indicate medians. Filled and open symbols indicate samples collected at earlier and later time points, respectively. Values below the lower limit of quantification were plotted at 1,000 copies/mL. Data were analyzed using the Kruskal–Wallis test followed by Dunn’s multiple comparisons test. Asterisks indicate significant differences versus controls. Blood samples at 10 dpi were available only from surviving mice. \**P* < 0.05 and \*\**P* < 0.01.

## Discussion

In this study, we showed that some cats that had recovered naturally from SFTS developed high neutralizing antibody titers and that their sera protected SFTSV-infected mice when administered early after infection. These findings provide proof of concept for further evaluation of passive immunotherapy as a treatment approach for feline SFTS, for which no established treatment or vaccine is currently available.

In our exploratory interspecies comparison, convalescent cats tended to have higher PRNT_80_ titers than convalescent human patients, although the difference was not statistically significant and titers varied among individuals (Figure 1). Because antibody responses likely differ between cats and humans, the comparison of PRNT_80_ titers between species should be interpreted cautiously. This trend may partly reflect the high blood viral RNA loads observed in cats with SFTS, which tend to exceed those reported in severe human cases; in turn, this may induce stronger humoral responses (Park et al., 2019; Gai et al., 2012). In addition, because SFTS predominantly affects older adults in Japan and lower IgG levels have been reported in older convalescent patients with SFTS, age-related differences in humoral responses may have influenced the antibody titers observed in human patients (Kobayashi et al., 2020; Li et al., 2023). Although differences in sampling time may have influenced the comparison of PRNT_80_ titers, studies in human patients with SFTS have shown that IgG and neutralizing antibodies peak several weeks to months after onset and can persist for years (Li et al., 2023; Wen et al., 2024). Further studies are needed to compare the quality, specificity, and effector functions of anti-SFTSV antibodies in cats and humans. Nevertheless, our findings suggest that naturally recovered cats may serve as a source of high-titer neutralizing antibodies, provided that donors are selected based on neutralizing antibody titers.

The therapeutic effect of convalescent cat serum was strongly influenced by timing of treatment. Administration at 2 or 3 dpi provided marked protection, whereas efficacy decreased when treatment was delayed to 4 dpi or later (Figures 2A and 3A). These outcomes are broadly consistent with previous studies of antiviral agents and monoclonal antibodies in SFTSV-infected mouse models (Kim et al., 2019; Tani et al., 2016), although direct comparisons are limited by differences in viral strains, inoculation doses, routes of administration, and mouse models.

Early treatment also reduced disease severity and viral replication. Although survival was similar after treatment at 2 or 3 dpi, body weight loss was less pronounced with earlier treatment (Figures 2B and 3B). Blood viral RNA levels were also lower when treatment was initiated at 2 or 3 dpi than when it was delayed to 4 dpi (Figure 2C). These findings suggest that convalescent cat serum becomes less effective after extensive viral replication has occurred. Because intraperitoneal antibody administration requires absorption from the peritoneal cavity before systemic exposure (Barrett et al., 1991), intravenous administration, which is more relevant to clinical use in cats, may achieve more rapid systemic exposure and should be evaluated in future studies.

Administration on two consecutive days did not substantially extend the therapeutic window for survival but improved suppression of late-phase viremia. Although mice treated at 2 and 3 dpi or 3 and 4 dpi were completely protected, delayed treatment showed limited efficacy. Nevertheless, viral RNA levels at 13 dpi were markedly lower after repeated administration than after single-dose treatment, suggesting that additional dosing may help control residual viremia. This effect could be clinically relevant because lower viremia may reduce disease severity and the risk of secondary transmission from infected cats to humans (Yamanaka et al., 2020; Mekata et al., 2023). The experiment using sera from individual convalescent cats further suggested that therapeutic efficacy may vary among donors, likely reflecting differences in neutralizing antibody titers (Figure 4).

A major limitation of this study is the use of AG129 mice, which lack type I and type II interferon receptors. This model was used because immunocompetent mice show only limited SFTSV replication and do not develop clinical disease (Liu et al., 2014). Although this model is useful for evaluating therapeutic effects under conditions of uncontrolled viral replication, it does not fully reproduce feline SFTS. Therefore, any survival benefit observed in AG129 mice should be interpreted as experimental proof of concept rather than as direct evidence of clinical efficacy in cats, particularly since SFTSV primarily targets macrophage/monocyte-lineage cells in AG129 mice but plasmablasts in cats (Matsuno et al., 2017; Sakai et al., 2021). Moreover, all control mice died in our experiments, whereas the reported case-fatality rate in naturally infected cats is approximately 60% (Matsuu et al., 2019; Ando et al., 2021). Therefore, the clinical value of this approach should ultimately be assessed through carefully documented clinical use in naturally infected cats. Although protection appeared to correlate with neutralizing antibody titers, we did not directly determine whether neutralizing antibodies alone accounted for the observed therapeutic effects. Because we did not perform IgG depletion, purified IgG transfer, or Fc-function analyses, contributions from Fc receptor-dependent antibody effector functions or other serum factors cannot be excluded.

Clinical application also raises safety and logistical issues. The serum volume used in mice was large on a body-weight basis and cannot be directly translated to clinical dosing in cats. The optimal dose, route, and frequency of administration in cats remain unknown, and feline blood products can cause severe transfusion reactions, which necessitates blood typing and cross-matching (Auer and Bell, 1983; McClosky et al., 2018). Implementation would also require SFTS-recovered donor cats with high neutralizing antibody titers that meet standard donor criteria, including negative testing for feline immunodeficiency virus and feline leukemia virus (Taylor et al., 2021).

In summary, some cats recovering from SFTS developed high neutralizing antibody titers, and high-titer convalescent cat serum protected SFTSV-infected mice when administered early after infection. These findings provide experimental proof of concept for convalescent serum-based passive immunotherapy for feline SFTS, but clinical studies in naturally infected cats are needed to determine its safety, efficacy, and practical applicability.

## Acknowledgments

We thank Saori Kusakari for technical assistance. We also thank the faculty and staff of the Department of Bio-resources, Frontier Sciences Research Center, University of Miyazaki, for their support with the introduction, breeding, and maintenance of AG129 mice. Finally, we are grateful to the cats and their owners for their cooperation in this study.

## CRediT authorship contribution statement

Conceptualization, H.M.; methodology, H.M.; validation, H.M.; formal analysis, H.M.; investigation, H.M. and M.Y.; resources, H.M., T.K., and T.O.; data curation, H.M. and M.Y.; writing—original draft preparation, H.M.; writing—review and editing, H.M., M.Y., T.K., and T.O.; visualization, H.M.; supervision, H.M.; project administration, H.M.; funding acquisition, H.M. All authors have read and approved the final manuscript.

## Data availability

The raw data supporting the findings of this study have been deposited in Figshare and will be made publicly available upon publication of the peer-reviewed version of this article: https://doi.org/10.6084/m9.figshare.32413089.

## Declaration of generative AI use

The authors used ChatGPT for language refinement, editorial assistance, and review of manuscript preparation materials. The tool was used to improve clarity, grammar, consistency with journal style, and the organization of manuscript-related text. The authors reviewed and edited all AI-assisted content, confirmed the originality and accuracy of the manuscript, and take full responsibility for the final content, including the accuracy of references.

